# Semen collection, short term storage, and cryopreservation in the Texas horned lizard (*Phrynosoma cornutum*)

**DOI:** 10.64898/2026.04.03.716302

**Authors:** Allison R. Julien, John A. Griffioen, Sean M. Perry, Robyn Doege, Isabella J. Burger, Diane Barber

## Abstract

As global reptile populations continue to decline, improving reproductive success in managed populations of listed species, such as *Phrynosoma cornutum* (the Texas horned lizard) has become increasingly critical for species survival. One understudied area of reproductive research in reptile species is gamete collection and storage, a crucial component for maintaining genetic diversity. In Texas, semen was collected from wild *P. cornutum* (n = 20) in June 2025. Semen collection was performed via electroejaculation (EEJ) under alfaxalone anesthesia. Prior to semen collection, snout-vent-lengths (SVL) and weights were recorded and testes measurements were taken using a portable ultrasound. Average sperm motility and concentration across all lizards was 83.7% and 85.7 × 10^6^ sperm/mL, respectively. While lizards with longer SVLs had higher sperm motility, weight and testis size did not affect sperm parameters. Samples were extended in INRA96 and divided for use in cold-storage longevity or cryopreservation trials. Samples under cold-storage conditions were assessed for motility daily for 10 days. Motility was not significantly reduced until 48 hours post-collection and maintained 19% motility at day 10. For cryopreservation, samples were diluted 1:1 in INRAFreeze cryopreservation media and frozen in liquid nitrogen, then immediately thawed. Average post-thaw sperm motility was 13.9%, with the highest post-thaw motility recorded at 38.2%. This is the first report of semen storage and cryopreservation in *Phrynosoma* and provides valuable insight into semen storage potential in reptile species.

## I. Introduction

With over 21% of reptile species today threatened with extinction (Cox et al. 2022; Farooq et al. 2024), the importance of maintaining assurance populations across zoological and conservation facilities has risen with an impetus on increasing reproductive output for species sustainability and reintroduction (Rahbek 1993; Clulow et al. 2022). However, due to their diverse reproductive strategies, complex gametic dynamics, and specific biotic and abiotic requirements (Angelini and Ghiara, 1984; Blackburn 1993; Shine 2003; Van Dyke 2015), breeding reptiles within managed populations can be challenging. Much of the research into the reproductive biology of reptile species has been largely focused on female endocrinology and reproductive function. Female follicular dynamics and the induction of ovulation and oviposition have been studied since the 1970s (Jones et al. 1976; Ewert and Legler 1978; Jones et al. 1988), with a focus on the treatment of dystocia and follicular stasis, also called egg binding, wherein oviposition is obstructed or impeded from a number of potential causes. Conversely, male reproductive research has been widely neglected. To date, few studies into the topics of semen collection, analysis, and storage have been conducted, and of these, fewer have related to application within lizard species (Clulow and Clulow 2016). This limited scale of study has slowed the development of reliable methodologies for techniques such as semen collection, sperm analysis, and preservation, impeding efforts for application in improving captive population management and reproductive output.

There are several benefits to sperm analysis within a breeding population. Primarily, the establishment of male reproductive health and fecundity is a critical component for breeding pairings and population management. This has been difficult to standardize across reptile species, however, due to the lack of study into sperm parameters. Establishing safe and successful protocols for semen collection has been a focal point for the development and improvement of sperm analysis (Perry 2021). Commonly in reptiles, semen collection is achieved through euthanasia of the animal and collection of semen from the vas deferens (Moshiri et al. 2014; Campbell et al. 2021), a method largely incompatible with managed populations of threatened and endangered species. As an alternative, manual collection of semen in living reptiles through massage has gained popularity in several species, including the crevice swift and Mexican horned lizard (e.g., Martínez-Torres et al. 2019) and the saltwater crocodile (Johnston et al. 2014; Johnston et al. 2017) and is the primary method for semen collection in snake species (Mattson et al. 2007; Sandfoss, Reichling, and Roberts 2023). Across species, however, successful collection and semen sample quality has been variable using this non-terminal method (Zimmerman and Mitchell 2017), limiting application of this methodology in breeding programs for non-snake species.

An alternative method for semen collection is electroejaculation (EEJ). Briefly, a rectal probe is inserted into the cloaca of the male, and an electrical current is used to stimulate the nerves surrounding the vas deferens, resulting in the expulsion of mature semen (Platz et al., 1980; Assumpção et al. 2017). The application of EEJ in reptile species has been validated across large-bodied species such as turtles (Platz et al. 1980; Zimmerman and Mitchell 2017), crocodilians (Assumpção et al. 2017), and the green iguana (Zimmerman, Mitchell, & Perry 2013; Perry et al. 2021). Due to commercial EEJ probe sizes and safety concerns, EEJ was not attempted in medium- and small-bodied lizards until 2018. In 2018, Juri et al. performed EEJ as a safe and successful method for semen collection in the medium-bodied spiny lava lizard (*Tropidurus spinulosus*) for a single timepoint. One year later, Martínez-Torres et al. published successful EEJ-induced semen collection in the crevice swift (*Sceloporus torquatus*). In the same year, the safety of repeated EEJ-induced semen collection was assessed by using health parameters in veiled chameleons (*Chamaeleo calyptratus*; Perry et al. 2019). Additionally, semen from veiled and panther chameleons (*Furcifer pardalis*) was repeatedly collected for a year to determine associations between semen production and hormonal cyclicity (Perry et al. 2023). Today, however, these methods have seen little expansion into other, smaller species, and their use is not widespread in conservation breeding programs despite repeated successes reported in the literature.

Due in part to the lack of established non-fatal semen collection and analysis protocols in reptiles, techniques for both short- and long-term semen storage remain one of the least developed assisted reproductive technologies (ART) for this taxon. Short-term refrigerated semen storage and cryopreservation are critical tools in conservation science and population management as both techniques allow for the salvage, transportation, and storage of valuable male genetics between and within managed populations (Clulow and Clulow 2016; Contreras et al. 2019; Perry et al. 2021; Anastas et al. 2023). Both techniques also have immediate utility in procedures such as artificial insemination (AI) and for the preservation of semen, ranging from short term (refrigeration) to perpetuity (cryopreservation). In order to safeguard reptile species against current and future threats of extinction and reduction in genetic diversity, it is imperative that these tools be developed for application in captive breeding programs.

The Texas horned lizard (*Phrynosoma cornutum*) is a medium-bodied lizard endemic to the south-central United States and northern Mexico (Ballinger 1974; Price 1990). In recent decades, populations have drastically declined in Texas (Price 1990; Williams, Rains and Hale 2019), leading to the species’ state listing as “threatened” in Texas. Assurance colonies and breeding populations have been established across several Texas zoological institutions as a means for mitigating further decline through the release of captive-bred offspring. Unfortunately, breeding and hatch rates across programs remain relatively low, hindering sustainability and reintroduction efforts. Due to the lack of established protocols for investigating and addressing reproductive failure in reptiles, efforts to improve fertility rates in managed care have been slow. Here, we aim to validate electroejaculation as a safe and reliable means of semen collection, determine cold-storage longevity of collected semen for future application, and investigate the current cryopreservation potential of collected semen from *P. cornutum*.

## II. Materials & Methods

### a. Animals

Wild adult *P. cornutum* (n= 20) were collected and sampled at the Matador Wildlife Management Area (MWMA) in Cottle County, Texas, in June of 2025 under Texas Park and Wildlife Permit #1091-455. Males were collected during road surveys and captured by hand. All animals were acclimated indoors overnight in plastic holding containers. Weights (g) and snout-vent length (SVL) measurements (mm) were taken for each male prior to anesthesia and semen collection. All research was conducted following review by Fort Worth Zoo’s Institutional Animal Care and Use Committee (IACUC).

### b. Anesthesia

Each individual was weighed and administered alfaxalone (Alfaxan® Multidose, Zoetis, Kalamazoo, MI 49007 USA) at a dose of 15-18 mg per kg body weight, rounded to the nearest hundredth by volume, in the left forelimb. Injections were administered intramuscularly, though some subcutaneous administration is expected given limited muscle mass of the forelimbs. Lizards were monitored via visual and reflex assessment for respirations and anesthetic depth. A Butterfly portable ultrasound (Butterfly iQ/iQ+™) linear probe compatible with Android was used to assess heart rate. Lizards were monitored for return of spontaneous reflexes and movements and were considered recovered when an immediate righting reflex returned.

### c. Testis measurements

Prior to electro-ejaculation, trans-abdominal ultrasound scans of each male were performed to assess testis size using the Butterfly portable ultrasound. Males were held in dorsal recumbency with their abdomens accessible. Ultrasound gel was applied to the abdomen of each individual and images were taken using the “nerve” setting at a gain (signal amplification) between 60% -70% at a depth of 3 cm. Two to five images were taken per individual, and testis length and diameter were measured using the Butterfly Network Cloud Software (**Figure 1**). Testes volumes (V) were then calculated using the following standardized equation from Sotos and Tokar (2012) and has been used repeatedly in reptiles:

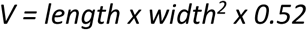

**Figure 1.**
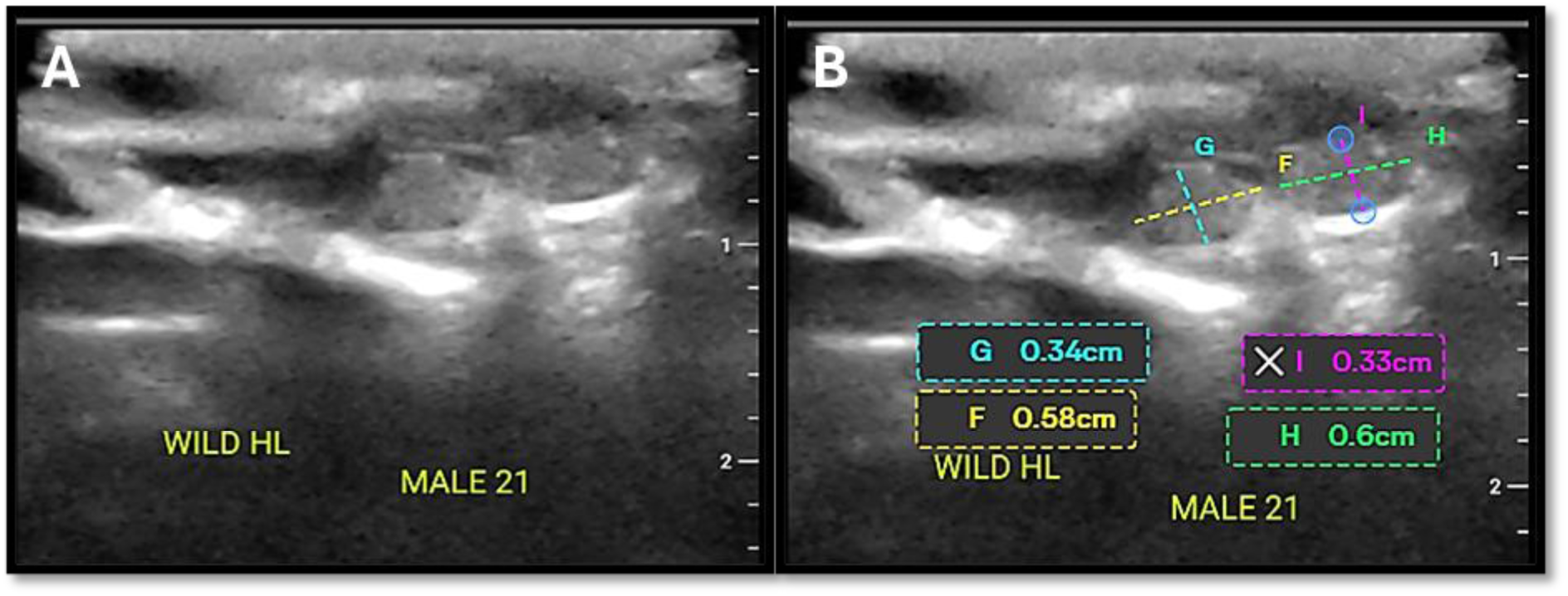
Ultrasound images of *Phrynosoma cornutum* testes (A) and testis measurements (B) under alfaxalone sedation

### d. Semen collection and analysis

Once a sufficient plane of sedation was achieved, males were laid on their dorsum and a 2 mm wide electroejaculation probe was inserted approximately 12 mm into the cloaca. Each male underwent a stimulation cycle consisting of 5 – 7 five-second discharges with a five second rest period in between each discharge. Stimulations were 3.0 - 5.0 V. After the stimulation cycle, the probe was removed from the cloaca and gentle pressure was applied to the abdomen in downward sweeping motion to draw semen down to the cloaca (**Figure 2**). Once semen was observed pooling at the cloacal opening, the semen was drawn up using a 20 µl pipette and placed into a bullet tube containing 100 µl of INRA96 semen extender (IMV Technologies) for storage at 4°C and subsequent analysis.

**Figure 2.**
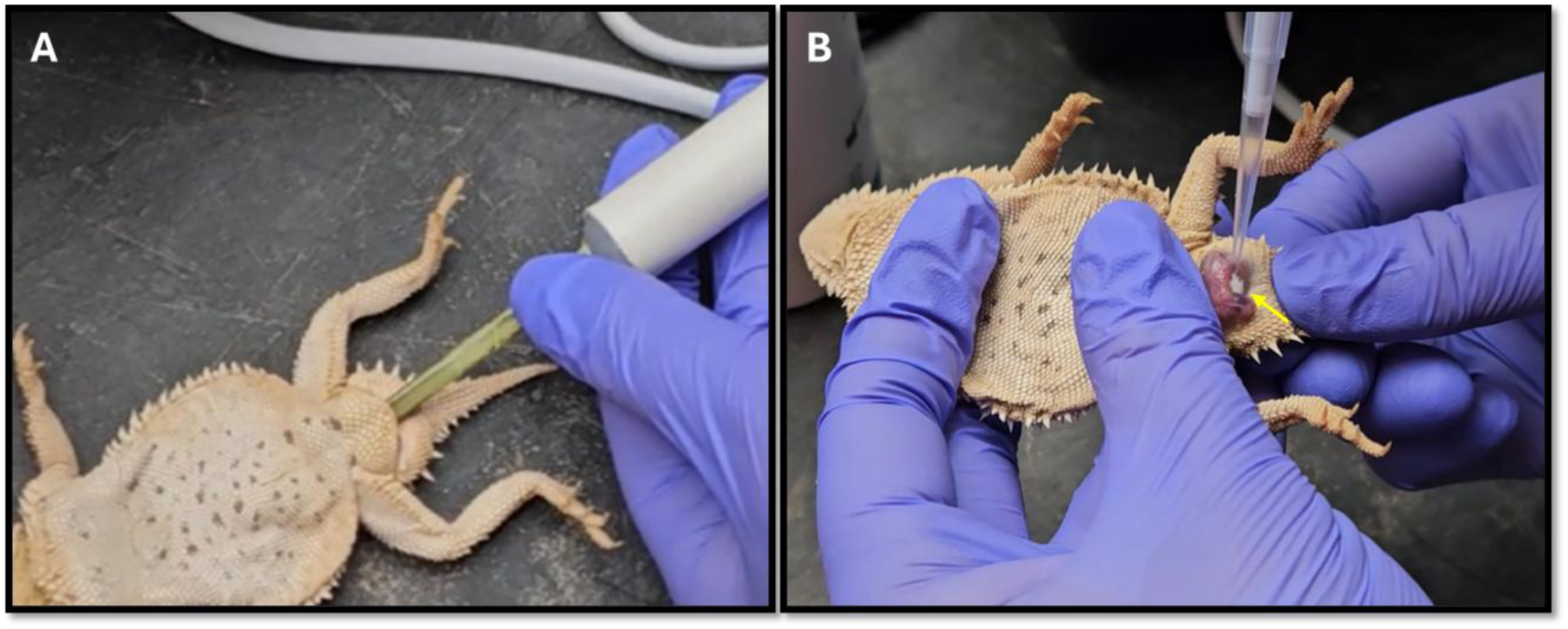
Insertion of electroejaculation probe into cloaca of male *Phrynosoma cornutum* (A) and semen collection using a pipette (B). Pooled semen at the cloacal opening is indicated with a yellow arrow

Following collection, semen volume was recorded before extender addition and sperm was analyzed for motility and concentration using an Olympus CX41 phase contrast microscope.

Sperm motility counts were conducted by counting individual sperm cells by hand using a multi-cell counter. Sperm motility was categorized as motile (M; sperm exhibiting flagellar movement but no forward movement), forward progressive motile (FPM; sperm exhibiting flagellar movement and progressing forward), and non-motile (NM; sperm exhibiting no flagellar movement). Motile and forward progressive motile values were added to describe total motility (TM). Sperm cells were counted until reaching 100 individual cells, giving the proportion of each cell motility type. Sperm concentration was counted on a Neubauer hemacytometer (Hausser Scientific) using a hand cell counter. Briefly, cells within the Neubauer ruling grids are counted. The number is then multiplied by 2500 to determine sperm per milliliter.

### e. Short term storage and cryopreservation

An additional 400 µl of INRA96 extender was added to each sample tube and gently mixed by pipetting the sample up and down. Samples were then divided in half with half reserved for testing cold storage longevity and half left for cryopreservation trials. To determine short term cold storage longevity, samples were stored in INRA96 semen extender at 4°C. A volume of 12 ul was removed daily for 10 days (240 hours) to assess changes in motility over storage time.

Cryopreservation trials occurred the same day as initial sample collection. Samples diluted in INRA96 were further diluted 1:1 in INRA Freeze (IMV Technologies) as a cryoprotectant media and similarly mixed using a pipette. Samples were then loaded into ¼ cc CASSOU semen straws (IMV Technologies) and frozen using a three-step slow freezing method. In step 1, loaded straws were left to equilibrate on an ice pack for 10 minutes. In step 2, equilibrated straws were placed into a Styrofoam freezing box containing liquid nitrogen at a level of 5 cm above the liquid nitrogen for an additional 10 minutes to allow slow vapor freezing. Straws were then plunged into liquid nitrogen.

Straws were thawed one at a time immediately following freezing. Straws were removed from liquid nitrogen and thawed in a 35°C water bath for approximately 3-5 seconds. Once removed from the water bath, straws were gently dried and emptied into Eppendorf tubes for analysis.

Samples were analyzed for post-thaw motility using methods outlined above. Post-thaw motility values were calculated as follows:

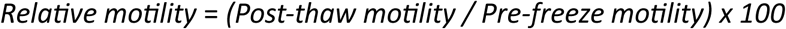

### f. Statistical analysis

To determine the effects of testes volume on sperm parameters, we ran three different analyses for total motility, forward progressive motility, and concentration. For total motility, we ran two linear models with total motility as the response variable and testes volume as a fixed effect. Because weight and SVL are correlated, we ran one model with SVL as a covariate and the other model with weight as a covariate. We then used AICc to determine if SVL or weight was a better predictor for total motility. Forward progressive motility and concentration did not meet assumptions of normality. Thus, we ran Spearman’s Rank Correlations for these two parameters, with forward progressive motility or concentration as the response, testes volume as the fixed effect and SVL, or weight as the covariate.

We were also interested in the effect of cold storage duration on motility. Therefore, we ran two linear mixed effects model with total motility or forward progressive motility as the response, time in cryostorage as the fixed effect, and male ID as a random effect to account for repeated measures. Lastly, we were interested in the effect of cryopreservation on motility. For total motility, we conducted a linear model with total motility as the response and freezing state (pre-freeze versus post-thaw) as the fixed effect. We initially included male ID as a random effect; however, it did not explain additional variation in the model and was removed. For forward progressive motility, we ran a linear mixed effects model with progressive motility as the response, freezing state as the fixed effect, and male ID as a random effect. We conducted all analyses in R (v 4.4.1). We tested for assumptions of normality using residual diagnostic plots.

## III. Results

### a. Effects of individual and testis size on sperm quality

Average male weight, snout-vent length, and testis volumes were 29.2 ± 0.9 g, 79.7 ± 0.9 mm, and 0.86 ± 0.01 mm^3^, respectively. Snout-vent length was a better predictor of total motility than weight (SVL AICc = 99.87; weight AICc: 110.56). Thus, we only discuss the results of the SVL analysis. SVL had a significant effect on total motility (F1,14 = 18.26, p = 0.0008), with larger individuals having more motile sperm. However, we did not find an effect of testes volume on total motility (F1,14 = 1.03, *p* = 0.33). For forward progressive motility, we found no effect of SVL (ρ(18) = 0.30, p = 0.21), weight (ρ(18) = 0.14; *p* = 0.56), or testes volume (ρ(18) = 0.17; *p* = 0.49). We also did not find an effect of these variables on sperm concentration (SVL: ρ(18) = -0.07, *p* = 0.79; weight: ρ(18) = 0.07, *p* = 0.79; testes volume: ρ(18) = 0.01; *p* = 0.97).

### b. Semen collection and analysis

Semen was successfully collected from all 20 sampled males. Ejaculate sizes were between 5 – 40 µl. Morphologically, sperm cells were characterized by long pointed heads with distinct distal droplets, likely signifying spermatozoa maturation, at the base of the head. Tails were approximately 3x the length of the heads (**Figure 3**). Average total sperm motility across all sperm samples was 85%, with an average forward progressive motility of 75.4%. The highest percentages of total motility and forward progressive motility were 96.0% and 87.0%, respectively, from separate individuals. The average sperm concentration across all sampled males was 81.5 × 10^6^ sperm/mL, with the highest concentration at 156.9 × 10^6^ sperm/mL.

**Figure 3.**
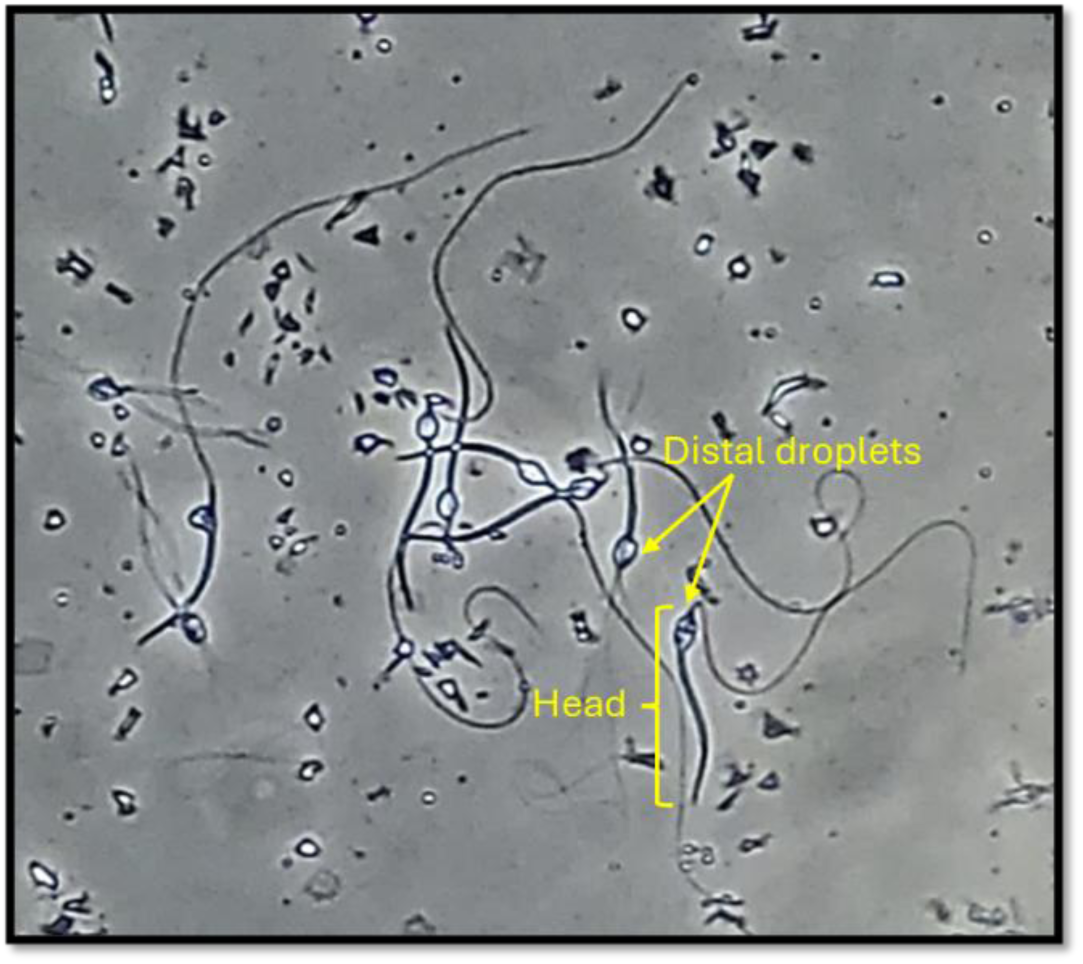
*Phrynosoma cornutum* sperm morphology indicating the heads of the spermatozoa (yellow bracket) and distal droplets on the heads (yellow arrows) following electroejaculation

### c. Short-term storage and cryopreservation

We found that sperm total motility and forward progressive motility significantly decreased with the duration of cold storage (TM: F10,172 = 17.58; *p* < 0.001; FPM: F10,172 = 16.70; *p* < 0.001), with values approaching zero after 240 hours. Total and forward progressive motility of sperm in cold storage were not significantly different from fresh sperm until after 48 hours (TM: p = 0.009; FPM: *p* = 0.009). Average total motility and forward progressive motility dropped below 50% after 192 hours and 144 hours, respectively (**Figure 4**).

**Figure 4.**
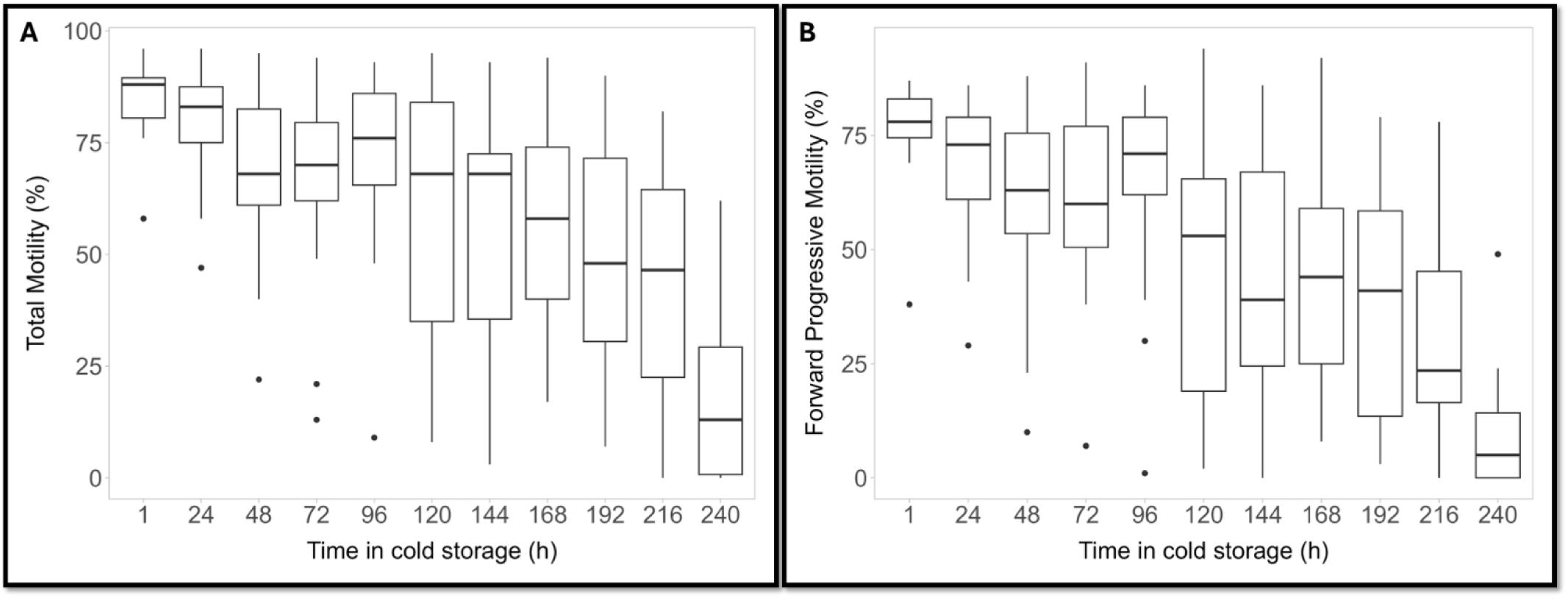
Average sperm total motility (A) and forward progressive motility (B) over time in cold storage following electroejaculation in the Texas horned lizard (*Phrynosoma cornutum*)

### d. Cryopreservation

Of the 20 males sampled in this study, semen samples from 19 males were selected for cryopreservation trials. One male was excluded due to low forward progressive sperm motility (52%). The average pre-freeze total motility across samples selected for freezing was 83.7%, with an average forward progressive motility of 75.9%. Out of the 19 males, only 1 male’s samples resulted in zero post-thaw motility, and 5 of the 19 males had over 20% relative post-thaw total motility. The overall average relative post-thaw total motility and forward progressive motility were 13.9% and 10.3% respectively. The highest reported post-thaw return for total motility reached 38.2%, with forward progressive motility having a maximum reported post-thaw motility of 44.6% (**Table 1**). Following post-thaw analysis, we found that both total (F_1,36_ = 547.56, *p* < 0.001) and forward progressive (F_1,18_ = 556.62, *p* < 0.001) post-thaw motility was significantly lower than pre-freeze motility (**Figure 5**).

**Figure 5.**
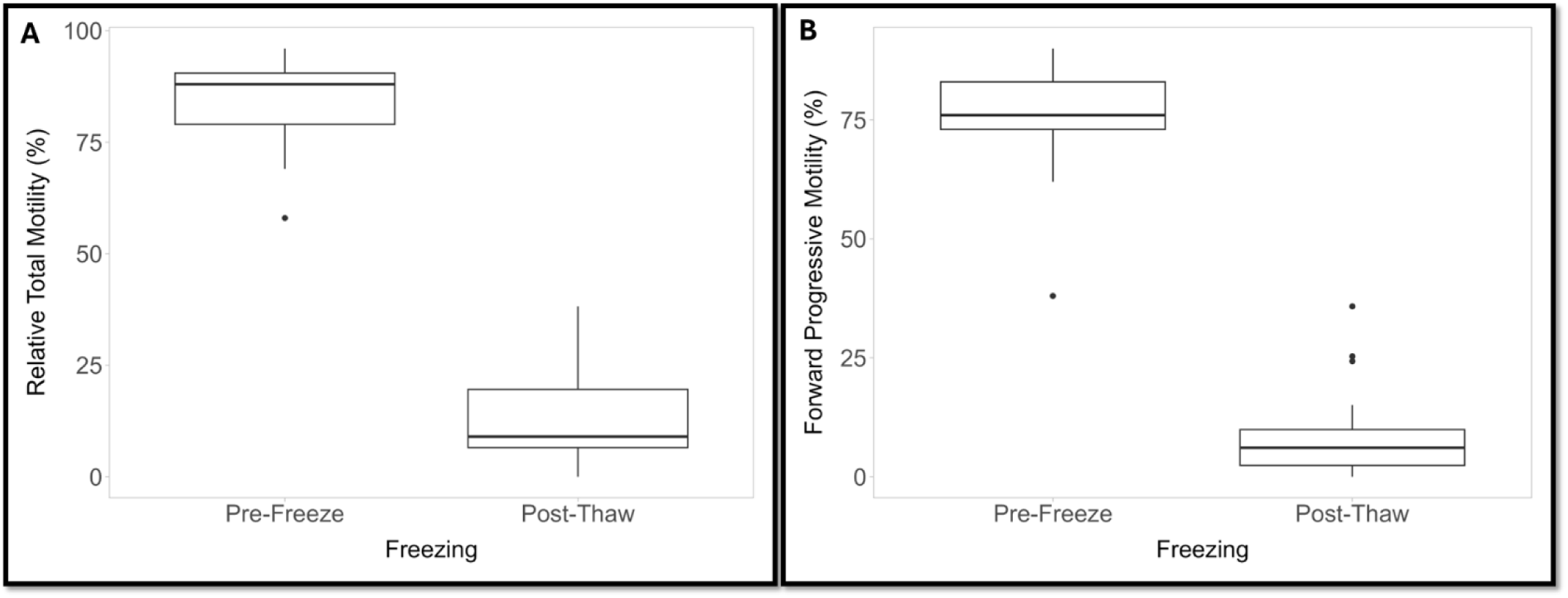
Average relative pre- and post- thaw total motility (A) and forward progressive motility (B) following cryopreservation trials Texas horned lizard (*Phrynosoma cornutum*)

**Table 1.** Male sperm concentration (sperm/mL) and motility values pre- and post-cryopreservation following electroejaculation in the Texas horned lizard (*Phrynosoma cornutum*). Values are shown as mean ± SEM

| Male | Sperm/mL<br>($\times 10^6$) | Forward Progressive Sperm<br>(%) | | Total Sperm Motility (%) | |
| --- | --- | --- | --- | --- | --- |
|  |  | Pre-Freeze | Post-Thaw | Pre-Freeze | Post-Thaw |
| 1 | 132.8 | 62 | 7.7 | 69.0 | 21.7 |
| 2 | 34.4 | 75 | 7.7 | 75.0 | 13.3 |
| 3 | 9.76 | 73 | 1.5 | 81.0 | 7.4 |
| 4 | 2.55 | 89 | 4.6 | 91.0 | 7.7 |
| 5 | 120 | 73 | 16.9 | 77.0 | 26.0 |
| 6 | 62.4 | 83 | 32.3 | 91.0 | 30.8 |
| 7 | 107.2 | 70 | 26.2 | 81.0 | 33.3 |
| 8 | 102.4 | 73 | 7.7 | 78.0 | 9.0 |
| 9 | 156.875 | 90 | 0.0 | 92.0 | 1.1 |
| 10 | 80 | 83 | 6.2 | 89.0 | 9.0 |
| 11 | 153.6 | 78 | 4.6 | 88.0 | 5.7 |
| 12 | 137.6 | 38 | 0.0 | 58.0 | 10.3 |
| 13 | 105.6 | 73 | 7.7 | 86.0 | 17.4 |
| 14 | 28.4 | 85 | 15.4 | 88.0 | 17.0 |
| 15 | 150.4 | 81 | 44.6 | 89.0 | 38.2 |
| 16 | 86.4 | 84 | 4.6 | 91.0 | 7.7 |
| 17 | 60.8 | 74 | 0.0 | 80.0 | 2.5 |
| 18 | 96 | 82 | 7.7 | 90.0 | 5.6 |
| 19 | 1 | 76 | 0.0 | 96.0 | 0.0 |
| Average | **85.7** | **75.9** | **10.3** | **83.7** | **13.9** |

## IV. Discussion

In the current study, electroejaculation (EEJ) resulted in the successful extraction of semen in all 20 of the sampled male *P. cornutum*. Furthermore, no deleterious effects were observed following observation over 2 hours post-treatment, which is consistent with the findings on the safety of EEJ in other lizards (Mitchel et al. 2015; Perry et al. 2021; Perry et al. 2022). Across all semen samples, average total motility and forward progressive motility was 85% and 75.4%, respectively. These are similar to the 78% percent motility reported in green iguanas (Zimmerman, Mitchell, & Perry 2013) and spiny lava lizards (Juri, Chiaraviglio, and Cardozo 2018). Sperm concentration (sperm/mL) was higher in the current study than in either of these studies; we report an average sperm concentration of 81.5 × 10^6^ sperm/mL, whereas the average concentrations reported in Zimmerman, Mitchell, & Perry (2013) and Juri et al. (2018) were 69 × 10^6^ and 2.1 × 10^6^ sperm/mL, respectively. This variation may be due to species-specific differences but may also be due to seasonality. The *P. cornutum* sampled in this study were sampled at the end of their historic breeding season (April – June; Ballinger 1974), while the experimental animals in Juri et al. (2018) were sampled outside of their breeding season (October – December; Cruz, Teisaire and Nieto 1997). The animals in Zimmerman, Mitchell, & Perry (2013) were sampled within the breeding season, but the authors note that recent mating may have reduced sperm quantity. Thus, in both studies the availability of mature spermatozoa may have been reduced compared to animals in the current study.

Following successful semen collection, we tested duration of both short-term and long-term sperm longevity. Short term storage, or refrigerated storage, refers to a cold storage technique wherein semen samples are stored in an extender and kept at low temperatures above freezing. Of the few reptile species wherein short-term storage has been attempted, sperm motility appears to demonstrate longevity spanning days to weeks (Molinia et al. 2010; Zimmerman, Mitchell, & Perry); however, results appear to be species-specific. Here, we report that sperm from wild *P. cornutum* remained motile for the entire 10 days (240 hours) of monitoring, with motility remaining above 50% for 8 of the 10 days. Total motility was reduced at an average rate of 7%/day, and forward progressive motility declined at a similar rate of 6.6%/day. The greatest drop in motility occurred for both total motility and forward progressive motility at day 10, wherein average motility decline dropped by over 20%. The longevity described herein provides evidence that horned lizard sperm can be stored and retain motility for up to 10 days, which is a large window during which horned lizard semen can be collected and utilized, making these results promising for the collection and transfer of semen from wild individuals or those that may be located in another breeding facility for use in artificial insemination attempts or cryopreservation.

Following successful short-term storage, we tested a protocol for semen cryopreservation in *P. cornutum*. Out of the 19 males from which samples were frozen, 18 retained motility following sperm cryopreservation, with an average total motility and forward progressive motility of 13.9% and 10.3%, respectively. This is the first instance of successful cryopreservation and post-thaw recovery of sperm in *P. cornutum*. Until 2016, there had been no published studies on lizard semen cryopreservation (Young et al. 2016). Since then, successful semen collection in lizards has expanded slightly to include the yellow-spotted monitor (*Varanus panoptes*; Campbell et al. 2020) and sceloporine lizard species (*Sceloporus aeneus*, *Sceloporus grammicus*, and *Sceloporus torquatus;* Sánchez-Rivera et al. 2022). Similar to short-term storage, protocols have also demonstrated species-specificity, limiting transferability and widespread protocol utilization. Our results are similar to those presented by Campbell et al. (2020), which tested post-thaw recovery of sperm from the yellow-spotted monitor in a DMSO-based cyroprotectant with and without post-thaw incubation in caffeine. Average post-thaw motility without caffeine incubation was reported as 18.1%, while post-thaw motility with caffeine was higher at 48%. Thus, future studies applying caffeine incubation post-thaw may be useful in *P. cornutum* to increase post-thaw sperm motility.

Finally, it is also important to note that the samples within the study were stored in the commercial semen media INRA96 and frozen in INRAFreeze media. INRA is a commercial semen extender for stallion sperm, containing milk micellar proteins, antibiotics, and fungicide. While the use of INRA96 as an extender for short-term storage was highly successful in this study, cryopreservation results were low for INRAFreeze as a cryoprotective media. However, the product protocol for INRAFreeze requires the addition of glycerol to the cryoprotective media. Reports in birds have noted that glycerol is a contraceptive (Hammerstedt and Graham 1992; Abouelezz et al. 2015); as one of the goals of cryopreservation is artificial insemination, we wanted to determine the success of the freezing media without the addition of glycerol. While INRAFreeze did not result in high post-thaw motility rates, it is currently a viable option for cryopreservation in *P. cornutum* that merits further study. As a semen extender, INRA96 is highly successful and readily accessible tool for short-term sperm storage in the species.

## V. Conclusion

Here, we present successful protocols for semen collection, analysis, and short-term storage in *P. cornutum*. These methods are poorly developed in reptiles, especially lizard species, and the development of successful protocols would drastically expand the horned lizard breeding strategies to include the transfer of semen from the wild or other institutions and/or long-term storage for future breeding. The sperm motility results following storage and cryopreservation presented here are the first to be reported in *P. cornutum* and are a valuable contribution to the literature surrounding lizard semen storage, cryopreservation, and to captive propagation of lizard species as a whole.

